# Ancient hybridization and adaptive introgression of an invadolysin gene in schistosome parasites

**DOI:** 10.1101/539353

**Authors:** Roy N. Platt, Marina McDew-White, Winka Le Clec’h, Frederic D. Chevalier, Fiona Allan, Aidan M. Emery, Amadou Garba, Amina A. Hamidou, Shaali M. Ame, Joanne P. Webster, David Rollinson, Bonnie L. Webster, Timothy J. C. Anderson

## Abstract

Introgression among parasite species has the potential to transfer traits of biomedical importance across species boundaries. The parasitic blood fluke *Schistosoma haematobium* causes urogenital schistosomiasis in humans across sub-Saharan Africa. Hybridization with other schistosome species is assumed to occur commonly, because genetic crosses between *S. haematobium* and livestock schistosomes, including S. *bovis,* can be staged in the laboratory, and sequencing of mtDNA and rDNA amplified from microscopic miracidia larvae frequently reveals markers from different species. However the frequency, direction, age and genomic consequences of hybridization are unknown. We hatched miracidia from eggs, and sequenced the exomes from 96 individual *S. haematobium* miracidia from infected patients from Niger and the Zanzibar archipelago. These data revealed no evidence for contemporary hybridization between *S. bovis* and *S. haematobium* in our samples. However, all Nigerien *S*. *haematobium* genomes sampled show hybrid ancestry, with 3.3-8.2% of their nuclear genomes derived from *S. bovis*, providing evidence of an ancient, introgression event that occurred at least 108-613 generations ago. Some *S. bovis* derived alleles have spread to high frequency or reached fixation and show strong signatures of directional selection; the strongest signal spans a single gene in the invadolysin gene family (Chr. 4). Our results suggest that *S. bovis*/*S. haematobium* hybridization occurs rarely, but demonstrate profound consequences of ancient introgression from a livestock parasite into the genome of *S. haematobium*, the most prevalent schistosome species infecting humans.

## Introduction

Introgressive hybridization occurs when hybrid offspring repeatedly backcross with one or both parental types, acting as a conduit for genetic exchange between species. Once present in the “new” genetic background, introgressed loci are broken up through recombination and exposed to selection. Deleterious alleles are purged while advantageous alleles may increase in frequency if they escape loss by drift. For example, genes impacting skin pigmentation (Vernot and Akey 2014) and immune response (Gittelman, *et al.* 2016) were transferred between Neanderthals and humans during the out-of-Africa migration(s). The gene underlying seasonal coat color changes in snowshoe hares are introgressed from jackrabbits (Jones, *et al.* 2018). Resistance to warfarin-containing-pesticides was transferred to mice from closely-related wild species (Song, *et al.* 2011), while resistance to insecticide-treated bed nets was transferred between *Anopheles gambiae* into *A. coluzzii* (Norris, *et al.* 2015). Introgression allows for complex phenotypic traits to be quickly introduced into a population over the course of a few generations allowing for rapid adaptation to new environments that may not be possible when relying on mutation or standing variation alone (Hedrick 2013).

Gene trspecificity or drug resistance to be transferred between ansfer occurs frequently between bacterial species through plasmid transfer or scavenging of environmental DNA, with profound ecological and biomedical consequences. Among parasitic organisms, hybridization and introgression is a concern since this provides a potential path for genes encoding biomedically important traits including virulence, host specificity or drug resistance to be transferred between species (King, *et al.* 2015). Schistosomes are dioecious trematodes that cause the debilitating disease schistosomiasis which infects more than 220 million people in 78 developing countries (World Health Organization 2019). *Schistosoma haematobium*, the focus of this paper, infects 112 million people (World Health Organization 2019) and is responsible for extensive morbidity including bladder cancer, genital schistosomiasis, and other pathologies associated with the urogenital tract. Schistosomiasis is associated with increased susceptibility to HIV and progression to AIDS (Colley, *et al.* 2014). While primarily confined to Africa and adjacent regions recent schistosomiasis outbreaks in Corsica caused by hybrids between *S. haematobium* and a closely related livestock species *S. bovis* have raised concern that this hybridization may have promoted range expansion and transmission potential (Boissier, *et al.* 2016; Kincaid-Smith, *et al.* 2018).

Interspecific laboratory crosses between members of *S. haematobium* species group, which contains nine closely-related species, have produced viable hybrid offspring (for examples see: Rollinson, *et al.* 1990; Bremond, *et al.* 1993; Tchuenté, *et al.* 1997; Southgatea, *et al.* 1998; Webster, *et al.* 2013) through to the F7 generation (Mutani, *et al.* 1985). In some cases, laboratory-reared, hybrid offspring have displayed hybrid vigor in the form of increased egg production (Mutani, *et al.* 1985), increased infectivity to intermediate and definitive hosts (Wright and Ross 1980), and expansion of suitable intermediate and definitive hosts (Huyse, *et al.* 2013). In natural populations, hybridization is diagnosed via a combination of pathology, egg morphology, and genetic identification via mitochondrial and nuclear discordance from single gene sequencing (reviewed in Leger and Webster 2017). In West Africa, parasites carrying ITS-rDNA from *S. haematobium*, but mtDNA from *S. bovis* are frequently observed and occasional parasites with ITS-rDNA copies from both species have been documented (Table 1). These results suggest hybridization, but the limited genotyping cannot fully distinguish whether this results from recent hybridization (F1 or F2 parasites), ancient hybridization and introgression, or incomplete sorting of ancestral lineages.

**Table 1.**
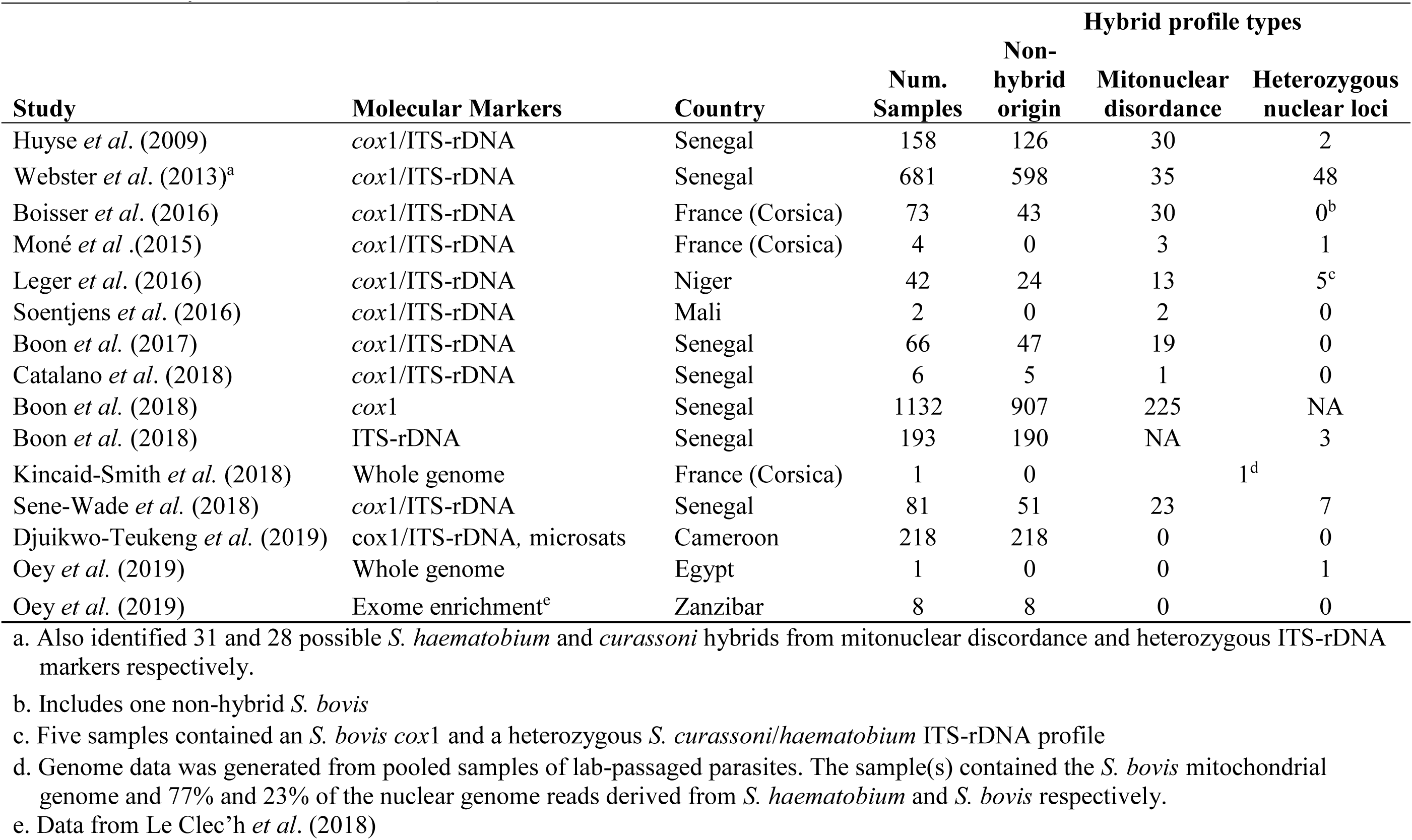
Summary of studies examining hybridization between *S. bovis* and *S. haematobium*.

Schistosomiasis ranks second only to malaria as a parasitic disease impacting human health. Yet unlike malaria for which thousands of parasite genomes have been sequenced (Pearson, *et al.* 2016; Amato, *et al.* 2018), population genomic analysis of schistosomes is in its infancy, mainly because adult worms live in the blood vessels of human hosts and cannot be easily sampled. In this study we examine hybridization and introgression in *S. haematobium*, leveraging a method that allows whole genome amplification followed by exome capture or whole genome sequencing from single miracidia larvae (Le Clec’h, *et al.* 2018). We sequenced exomes from parasite populations from both East Africa (Zanzibar) and West Africa (Niger). We used these data to examine the distribution of haplotype blocks containing *S. bovis* alleles within the *S. haematobium* autosomal genome to determine (A) whether hybridization is ancient or ongoing, (B) to evaluate evidence for introgression, and (C) to determine whether introgressed alleles show signatures of selection.

## Results

### Parasite samples, exome capture and sequencing

We used individual archived *S. haematobium* miracidia larvae from Niger and Zanzibar (Tanzania) fixed on Whatman FTA cards from the <u>S</u>chistosome <u>C</u>ollection <u>a</u>t the <u>N</u>atural History Museum (SCAN; Emery, *et al.* 2012). To minimize any impact of within host population structure (Steinauer, *et al.* 2013), we used a single miracidium from each of 96 different *S. haematobium* infected people (*n*_Zanzibar_=48, *n*_Niger_ = 48) for this study (Table S1). The Zanzibar samples were collected in 2011 and came from 26 locations on Unguja and Pemba islands spaced up to 160.9 km apart (Figure 1). The Niger samples were collected in 2013 from school-aged children from 10 locations along the Niger River located up to 125 km apart. We sequenced exomes from 88 miracidia from Niger (n=48) and Zanzibar (n=40) using whole genome amplification and exome capture (Le Clec’h, *et al.* 2018). The exome capture probe set was designed using the *S. haematobium* reference sequence contained 156,004,120bp RNA baits accounting for 94% of the exome length (15,002,706 bp/15,895,612 bp) in the nuclear genome as well as 67 RNA baits targeting the mitochondrial genome. Sequence coverage averaged 40.75x (range: 5.58-98.27; Table S1) across the 15Mb region targeted by our baits after excluding a single sample outlier sample that did not generate much sequence data. We combined our data with eight exome sequences from Zanzibar that have been reported previously for a total of 47 from Zanzibar (Le Clec’h, *et al.* 2018) and 48 from Niger.

**Figure 1.**
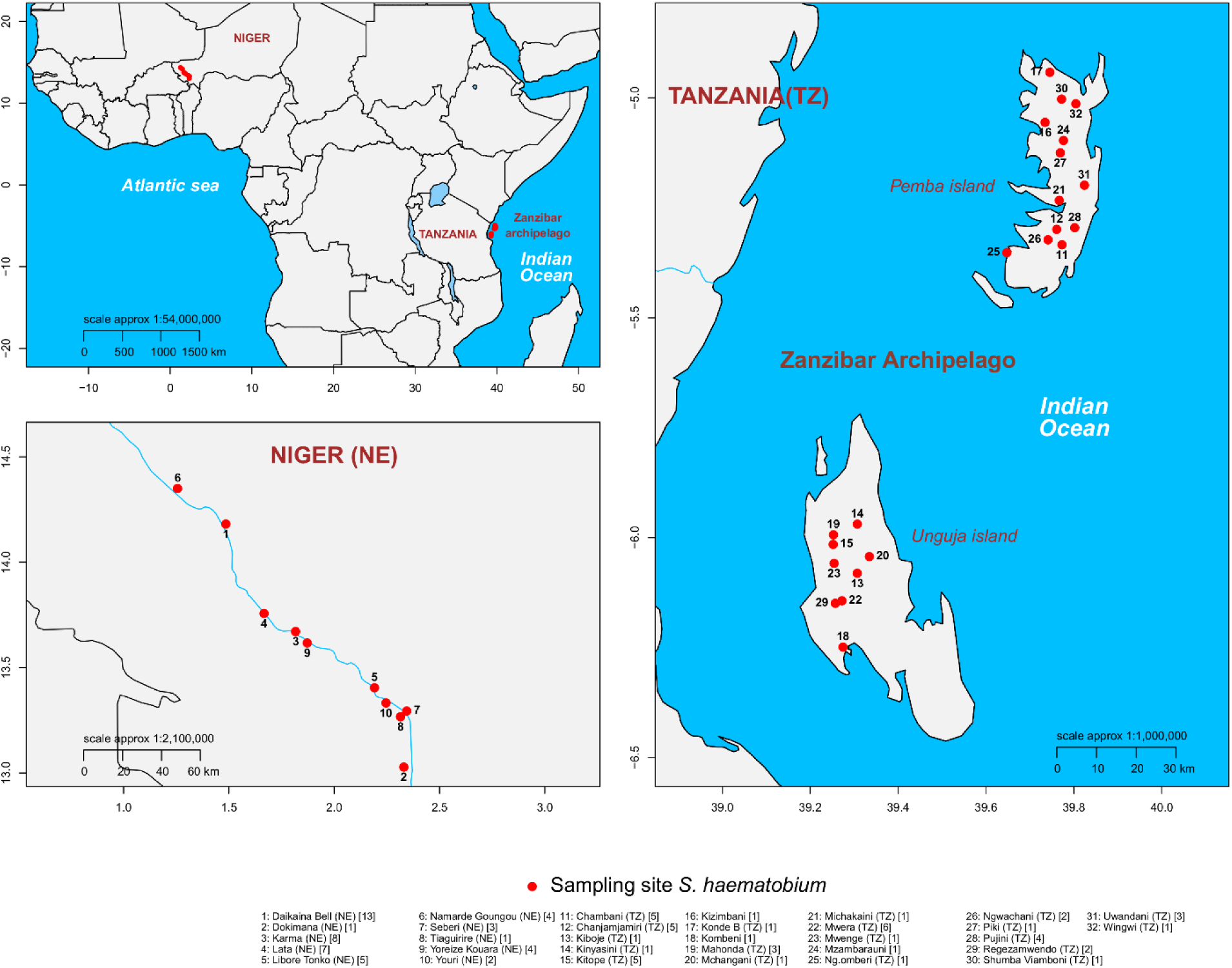
Collection localities. *S. haematobium* samples were collected in Niger and on the Zanzibar Archipelago off the coast of Tanzania.

### Mitochondrial variation

Our initial attempts to genotype mitochondrial loci contained within the exome probe set failed in 31 of the Nigerien S. *haematobium* samples due to poor representation of mitochondrial reads in the sequence data, despite high rates of capture from nuclear loci (Figure 2). We therefore sequenced whole genomes from a subset (n=12) of parasites to an average coverage of 17.8x and *de novo* assembled the mitochondrial genomes for each individual, while we genotyped *cox1* in other samples by Sanger sequencing. We found that the samples that failed mitochondrial probe capture contained a *S. bovis*-derived mitochondrial haplotype that was 15-18.05% divergent from that of *S. haematobium* and therefore poorly captured by our exome capture baits, designed from the *S. haematobium* reference sequence (Bi, *et al.* 2012). In all, 65% (*n*=31) of the Nigerien *S. haematobium* miracidia examined had a *S. bovis* mtDNA, similar to other West African *S. haematobium* populations (Huyse, *et al.* 2009; Webster, *et al.* 2013). In contrast, *S. bovis* mtDNA was not present in the Zanzibari *S. haematobium* population.

**Figure 2.**
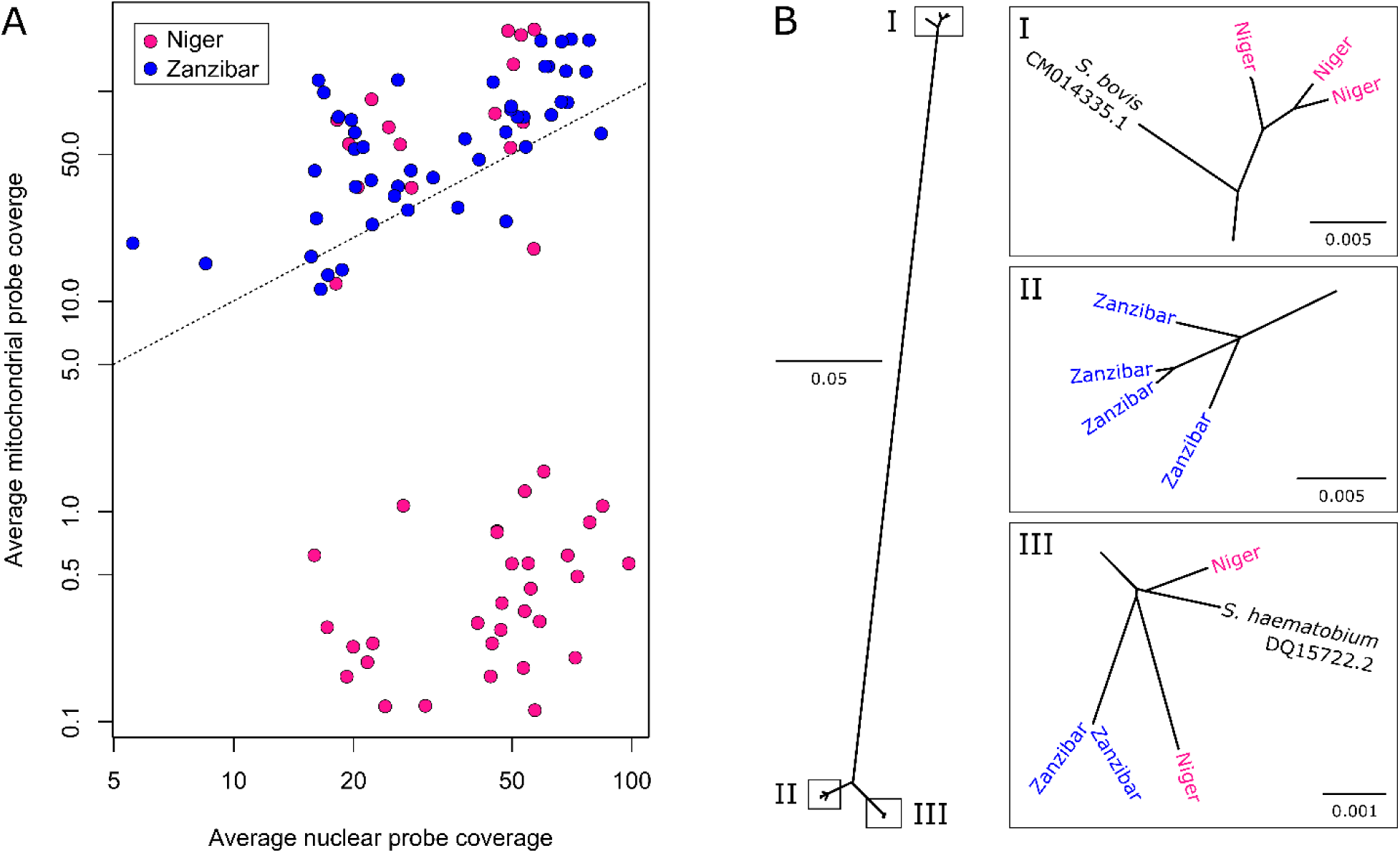
Mitochondrial sampling and phylogeny. (A) The mitochondrial genome was sequenced to high coverage despite large variations in the number of reads generated for most samples. For 31 Nigerien samples, mitochondrial reads were at very low frequency regardless of sequencing depth. This was attributed to the inefficient capture of the highly diverged mitochondria in these 31 Nigerien samples. (B) Maximum likelihood phylogeny of 12 mitochondrial genomes from *S. haematobium* and *S. bovis* and *haematobium* reference sequences. Three Nigerien *S. haematobium* samples are related to *S. bovis* indicating mitochondrial introgression from *S. bovis* into the Nigerien *S. haematobium* population.

### Exome variation

The presence of *S. bovis* mtDNA in Nigerien *S. haematobium* could result from contemporary hybridization or past introgression events. To differentiate between these two scenarios, we examined-autosomal, exonic (single nucleotide polymorphisms) SNPs recovered by the exome capture probes (Chevalier, *et al.* 2014; Le Clec’h, *et al.* 2018). In addition to our 96 *S. haematobium* samples we used sequence data from previous studies (Young, *et al.* 2012; Coghlan, *et al.* 2019) of six closely-related species in the *S. haematobium* species group (*S. bovis*, *S.* c*urassoni, S. intercalatum, S. guineensis, S. mattheei,* and *S. margrebowiei*). The genome assembly of *S. haematobium* is highly fragmented in thousands of scaffolds compared to the *S. mansoni* assembly, which comprises large chromosomal length scaffolds. However, the genomes of *S. haematobium* and *S. mansoni* are ≥89.4% syntenic (Young, *et al.* 2012). We aligned the *S. haematobium* and *S. mansoni* assemblies to convert SNP coordinates from their position on *S. haematobium* contigs to the corresponding position on *S. mansoni* chromosomes to take advantage of the more contiguous assembly and gene annotations available for *S. mansoni* (Berriman, *et al.* 2009; Protasio, *et al.* 2012). We verified local synteny of the transposed SNP coordinates by comparing linkage disequilibrium (LD) in 1Mb windows between the two sets of SNP positions (Figure S1). We recovered 370,770 autosomal, exonic SNPs for all samples including outgroups after genotyping, filtering and scaffolding along the *S. mansoni* assembly. These included 185,601 SNPs segregating in Niger and Zanzibar *S. haematobium* populations, of which 35,102 were common (MAF > 0.05). We found higher SNP diversity (paired t-test; df = 20,825; *p* = 2.2 × 10^−16^) among common alleles in Niger (mean π = 7.97 × 10^−5^; interquartile range = 6.3 × 10^−5^) than in Zanzibar (mean π = 2.89 × 10^−5^; interquartile range 2.3 × 10^−5^)

### Quantifying and dating admixture

We removed SNPs showing strong linkage disequilibrium (LD) to generate a reduced SNP subset for investigating population structure. A principle component analysis (PCA) from 5,882 unlinked, autosomal SNPS (MAF > 0.05) and all *Schistosoma* samples clearly differentiated *S. haematobium* from all other species along the first two PCs which accounted for 48.9% of genotypic variation observed (Figure 3A). The lack of intermediate genotypes between *S. haematobium* and other schistosome species suggests that we did not have early generation hybrids within our samples. We used ADMIXTURE (Alexander, *et al.* 2009) to assign ancestry proportions to the *S. haematobium*, *bovis*, and *curassoni* samples. *S. curassoni* was included due to its close phylogenetic affiliation with *S. bovis* (Lockyer, *et al.* 2003). Three distinct ancestry components were identified corresponding to *S. bovis*/*curassoni*, Nigerien, and Zanzibari *S. haematobium*. The *S. bovis*/*curassoni* ancestry component was present in 16 of the 48 Nigerien *S. haematobium* indicating low levels of potential admixture (0.1 < *Q* > 2.7; Figure 3B), while no admixture was identified in the Zanzibari *S. haematobium*.

**Figure 3.**
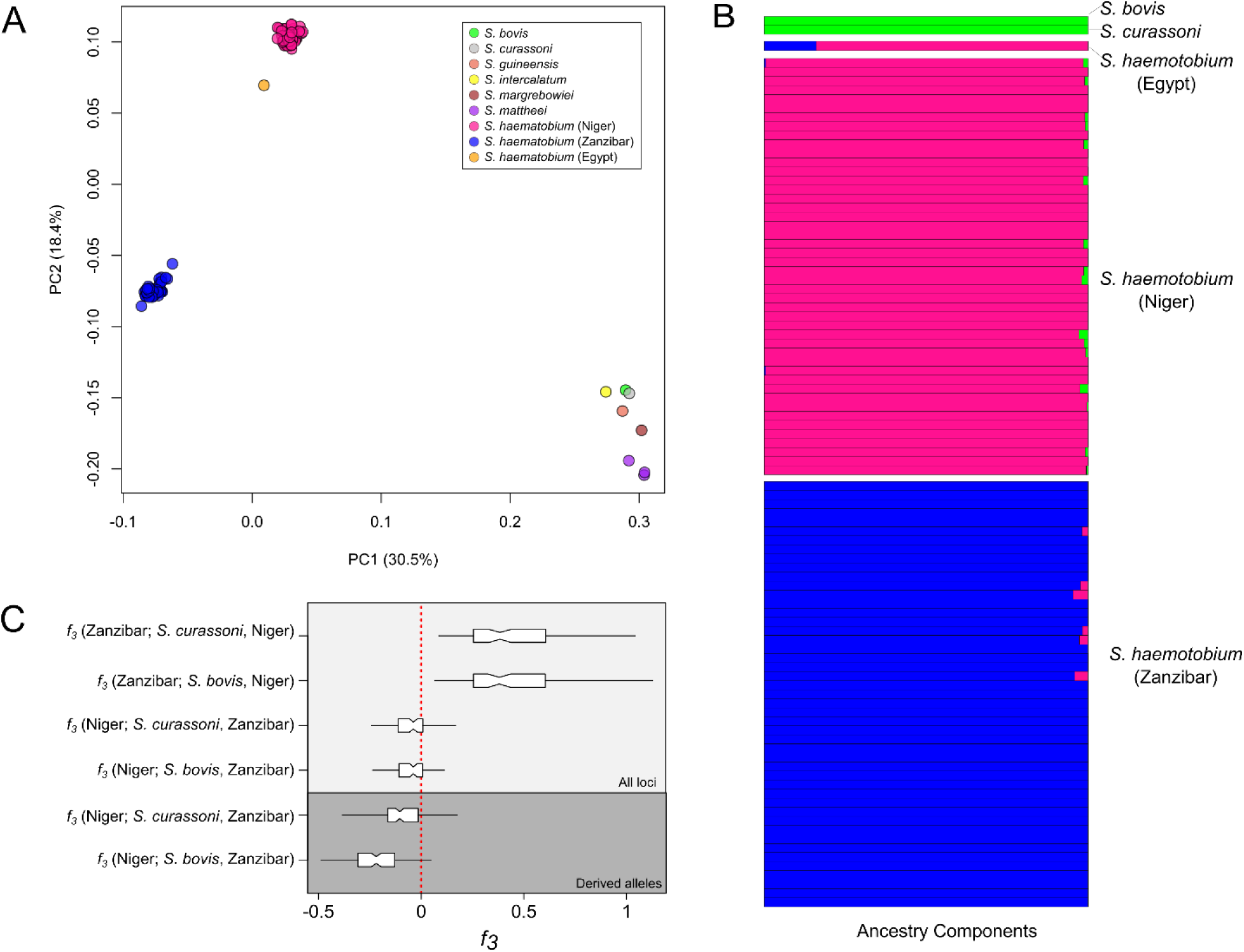
Population structure in *Schistosoma haematobium*. (A) PCA plot shows clear distinction between the two *S. haematobium* populations and the rest of the species examined: we see no evidence for recent hybridization (B) Quantification of population structure with ADMIXTURE showed low levels of admixture between *S. haematobium* populations and *S. bovis*/*curassoni*. (C) The three-population statistic (f3) was used to formally test admixture between each of the *S*. *haematobium* populations, *S*. *bovis*, and *S*. *curassoni*. When testing f3 (test; A, B), a negative result indicates that the test group is an admixed population from A and B. To differentiate between introgression from *S*. *bovis* and *curassoni* in Nigerien *S*. *haematobium*) we recalculated f3 using only derived alleles from *S. bovis* or *curassoni*. These results rule out contemporary hybridization between populations of *S. haematobium* and *S*. *bovis* or *curassoni*, and indicate that low levels of introgression are restricted to the Nigerien *S*. *haematobium* population.

We used the three-population test (*F_3_*) to determine whether *S. bovis* and/or *S. curassoni* were sources of introgressed alleles in the Nigerien *S. haematobium* population (Reich, *et al.* 2009). The *F_3_* test determines whether a target population is admixed from two source populations by calculating the product of differences in allele frequencies between the target and each source population. A negative *F_3_* indicates the target is an admixed population from the two sources. We used the Zanzibar S*. haematobium* population as a representative of the “source” *S. haematobium* population and alternated *S. bovis* and *S. curassoni* as the second source. *F_3_* results when testing for admixture from *S. bovis* (*F_3_* = −0.05, SE = 0.001, Z = −5.73) and *S. curassoni* (*F_3_* = −0.05, SE = 0.01, Z = 5.28) were negative indicating admixture from both species in the Nigerien *S. haematobium* population (Figure 3C). *S. bovis* and *curassoni* are very closely related. The admixture signature from *F_3_* could be driven by shared variation from the *S. bovis* and *curassoni* common ancestor. We recalculated *F_3_* from, but only using *S. bovis* or *curassoni* derived alleles minimize the signal from shared sites. These data provided evidence for admixture between *S. haematobium* and *S. bovis* (*F_3_* = −0.23, SE = 0.0129, Z = −18.18) but failed to identify admixture from *S. curassoni* (*F_3_* = −0.03, SE = 0.025, Z = 1.36). Incomplete lineage sorting provides a possible explanation when closely related alleles are found within different species. We confirmed that the *S. bovis* alleles in the Nigerien *S. haematobium* samples were the result of introgression as opposed to incomplete lineage sorting using Patterson’s *D* (D = 0.144; SE = 0.04; Patterson, *et al.* 2012). From these results, we infer that *S. bovis* alleles in Nigerien *S. haematobium* are the result of admixture.

We used PCAdmix (Brisbin, *et al.* 2012) to identify locally introgressed alleles in the Nigerien *S*. *haematobium* population. Ancestry was assigned to phased haplotype blocks by comparing them to a reference panel of *S. bovis* and Zanzibari *S. haematobium*. Within each Nigerien *S. haematobium* miracidium 3.3-8.1% (*x̅*= 5.2%; Figure 4) of alleles were identified as *S. bovis* haplotypes. In contrast, we observed close to zero *S. bovis* ancestry in Zanzibari miracidia used as a control (0.004 − 0.2%). Combined, these results indicate at least one introgression event between Nigerien *S. haematobium* and *S. bovis.* In comparison, Zanzibari *S. haematobium* do not contain *S. bovis* alleles.

**Figure 4.**
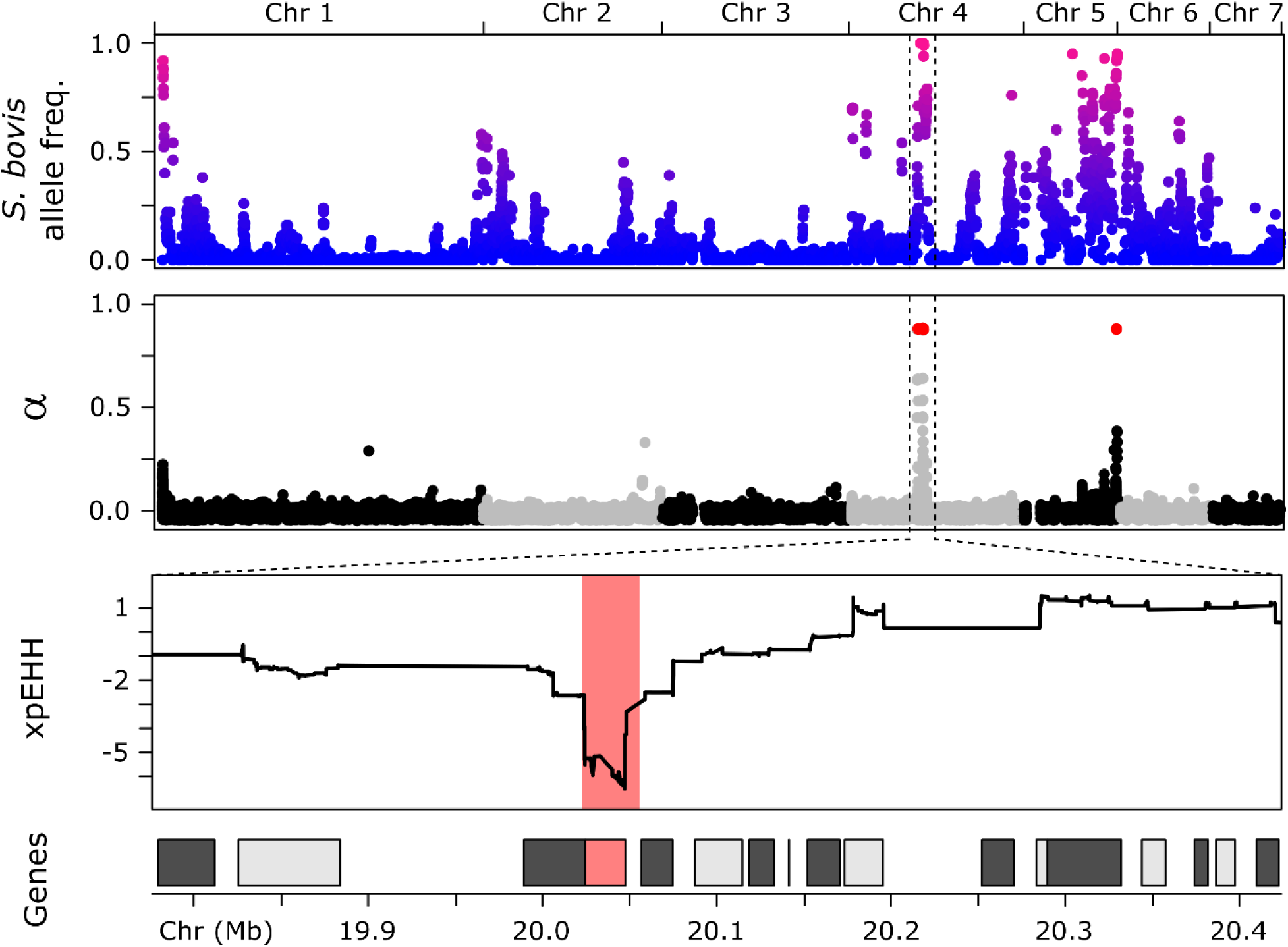
Selection on introgressed alleles in Nigerien *S. haematobium*. We aligned plots showing (a) proportion of *S. bovis* ancestry (b) genetic differentiation (FST) between Niger and Zanzibari samples (c) the α statistic from BayeScan and (d) xpEHH. *S. bovis* alleles have reached high frequencies in the Nigerien *S. haematobium* population. Introgressed alleles under directional selection were identified using allele frequency differences (α; alpha) between populations and using regions of extended homozygosity (xpEHH). Each test identified multiple regions under selection, but only one region was identified by both approaches. This region on Chr 4 spanned a single gene (Smp_120703, invadolysin) at which *S. bovis* alleles are approaching fixation in Nigerien *S*. *haematobium*.

Recombination breaks down introgressed segments over time. As a result, the length of introgressed haplotype blocks can be used to date time since admixture in number of recombination events or generations (Schumer, *et al.* 2016). Using this method we identified 5,649 *S. bovis* haplotype blocks in Nigerien *S. haematobium* averaging 0.5 cM in length and estimated admixture to have occurred ∼240.6 generations ago (min = 107.8 max = 612.5; Figure 5). Estimates of *Schistosoma* generation time vary greatly: while adult worms can live up to 37 years (Chabasse, *et al.* 1985), the minimum time for eggs to mature into adult worms is estimated to be ∼85 days in a laboratory setting (Loker 1983; Crellen, *et al.* 2016). Additional factors including water temperature (Lawson and Wilson 1980), host availability, and drug treatment will further influence average generation times. Assuming a ∼1 year generation time (King, *et al.* 2000) the hybridization event leading to introgression occurred ∼240.6 years ago (min = 107.8 max = 612.5). Importantly, while all 48 Nigerien miracidia contain introgressed *S. bovis* DNA resulting from ancient hybridization, we observed no F1 or early generation hybrids. Given 0/48 early generation hybrids detected in our population sample, we infer that the population frequency is between (0-7.4%; 95% binomial confidence interval; Clopper and Pearson 1934)

**Figure 5.**
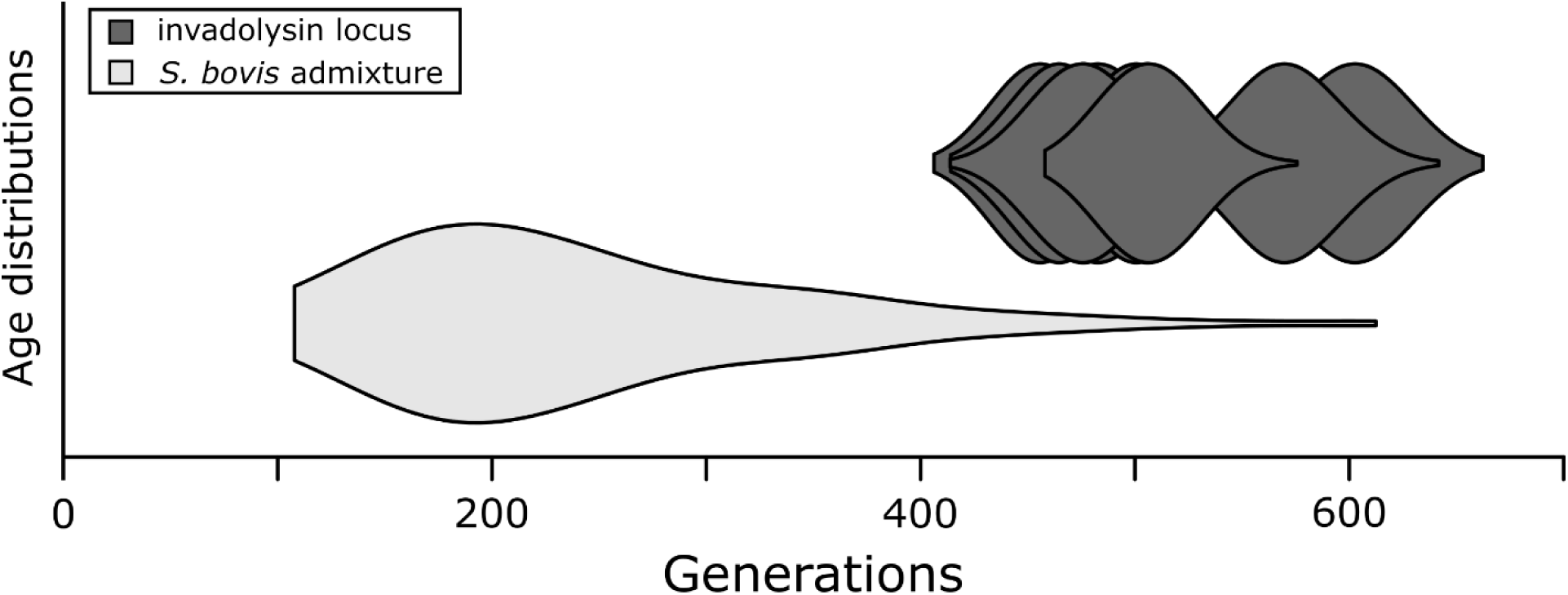
Time since admixture. Time since admixture was estimated using introgressed haplotype block length using methods described in Schumer, *et al.* (2016). The age of the invadolysin (Smp_127030) locus specifically (nruns = 10), was aged using startmrca (Smith, *et al.* 2018). Age estimates for both methods overlap, but the distribution of time since introgression tends to be younger than the estimated age of the invadolysin (Smp_127030) locus. Estimates of time since introgression could be influenced by exome data, which is limited in its ability to identity small haplotype blocks and result in conservative, younger age estimates.

### Selection on introgressed alleles

Since the admixture event(s) some *S. bovis* alleles are approaching or have become fixed in the Nigerien *S. haematobium* population (Figure 4). We used two complementary methods to examine directional selection in the Zanzibari and Nigerien *S. haematobium* samples. First, BayeScan (Foll and Gaggiotti 2008) was used to quantify selection (α) as a function of allele frequency differences at individual SNPs. Then large haplotype blocks under directional selection were identified using cross-population extended haplotype homozygosity (xpEHH; fig S2; Sabeti, *et al.* 2007). BayeScan and xpEHH identified 2 and 17 regions under directional selection between the Zanzibari and Nigerien *S. haematobium* populations. The strongest signals of selection from both analyses (α > 1.75; xpEHH ≤ −3) occurred at a 23Kb locus on chromosome 4 (Chr4:20,023,951-20,047,325; Figure 4), which also overlapped the region with the highest frequency of introgressed *S. bovis* alleles in the Nigerien *S. haematobium* population.

Selection of introgressed *S. bovis* alleles would be expected to purge variation in the vicinity of these alleles (Smith and Haigh 1974; Stephan 2019). We therefore plotted the log ratio of diversity (π) in Niger and Zanzibar samples, to identify genome regions showing exceptionally low diversity in Niger. The 23Kb locus genome region on Chr. 4 falls within the bottom 0.2% of values genome wide, consistent with the expectations of a selective sweep (Figure 6).

**Figure 6.**
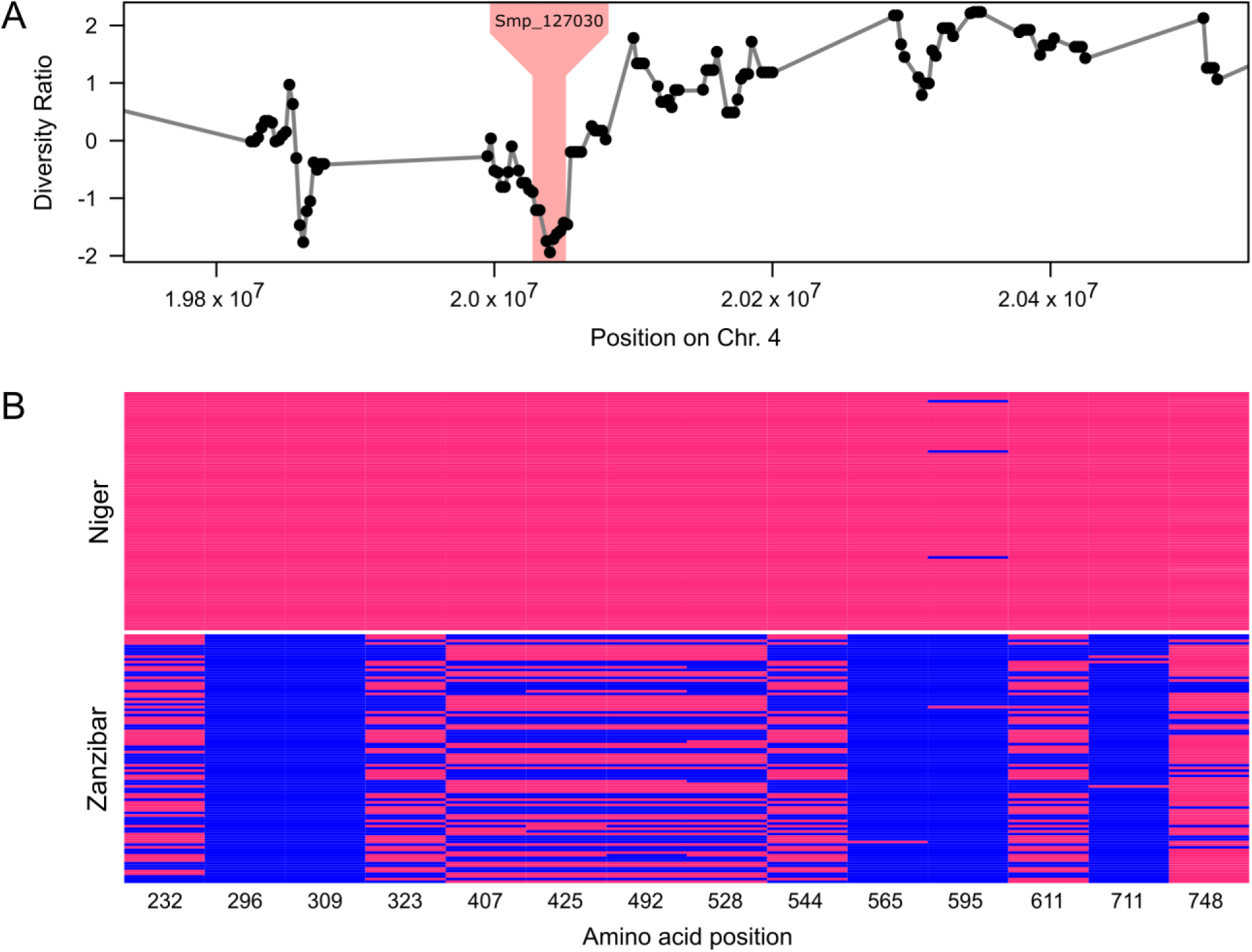
Reduced diversity at the invadolysin (Smp_127030) locus in the Nigerien *S*. *haematobium* population. (A) We compared differences in nucleotide diversity (π) across the Nigerien and Zanzibari *S*. *haematobium* populations calculated as log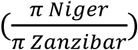. The region containing invadolysin (shaded) shows reduced variation (in the bottom 0.2% genome-wide) in the Nigerien *S. haematobium* population. (B) We identified 44 SNPs in Smp_127030, of which 14 resulted in non-synonymous changes (shown) and were also shared by more than one individual.

To confirm that this locus was the product of an introgression event rather than the result of selection on standing variation, we calculated the time to the common ancestor of the 23 Kb locus in the Nigerien haplotypes relative to the surrounding genomic regions using startmrca (Smith, *et al.* 2018). Using mutation (8.1*e*^-9^/bp/generation) and recombination rates (244.2 kb/cM), reported for *S. mansoni* (Criscione, *et al.* 2009; Crellen, *et al.* 2016), we estimated the time since divergence of the 23 Kb target region in the Nigerien haplotypes to be 476 generations ago (437.9-616.6; 95% CI; Figure 5), well within the range of our previous estimates of introgression from the haplotype block length analysis.

### Characteristics of the introgressed invadolysin gene

The 23 KB, introgressed region spans a single gene alternatively referred to as Smp_127030 (*S. mansoni* protein ID), MS3_06416, (*S. haematobium* protein ID), leishmanolysin, or invadolysin. Since we are using *S. mansoni* genome annotations and chromosomal coordinates we will refer to the invadolysin gene by its *S. mansoni* protein ID; Smp_127030. Within the invadolysin gene, we found 23 SNPs coding for 14 nonsynonymous changes (excluding singletons) two of which show fixed differences between Zanzibar and Niger populations (Figure 6). We were able to place six of the 14 nonsynonymous amino acid changes on a protein structure of Smp_127030, modeled from the *Drosophila* invadolysin (Protein DB: d1lmla). All were in peripheral positions on the protein (Figure 7) with little impact on the inferred protein structure (Kelley, *et al.* 2015).

**Figure 7.**
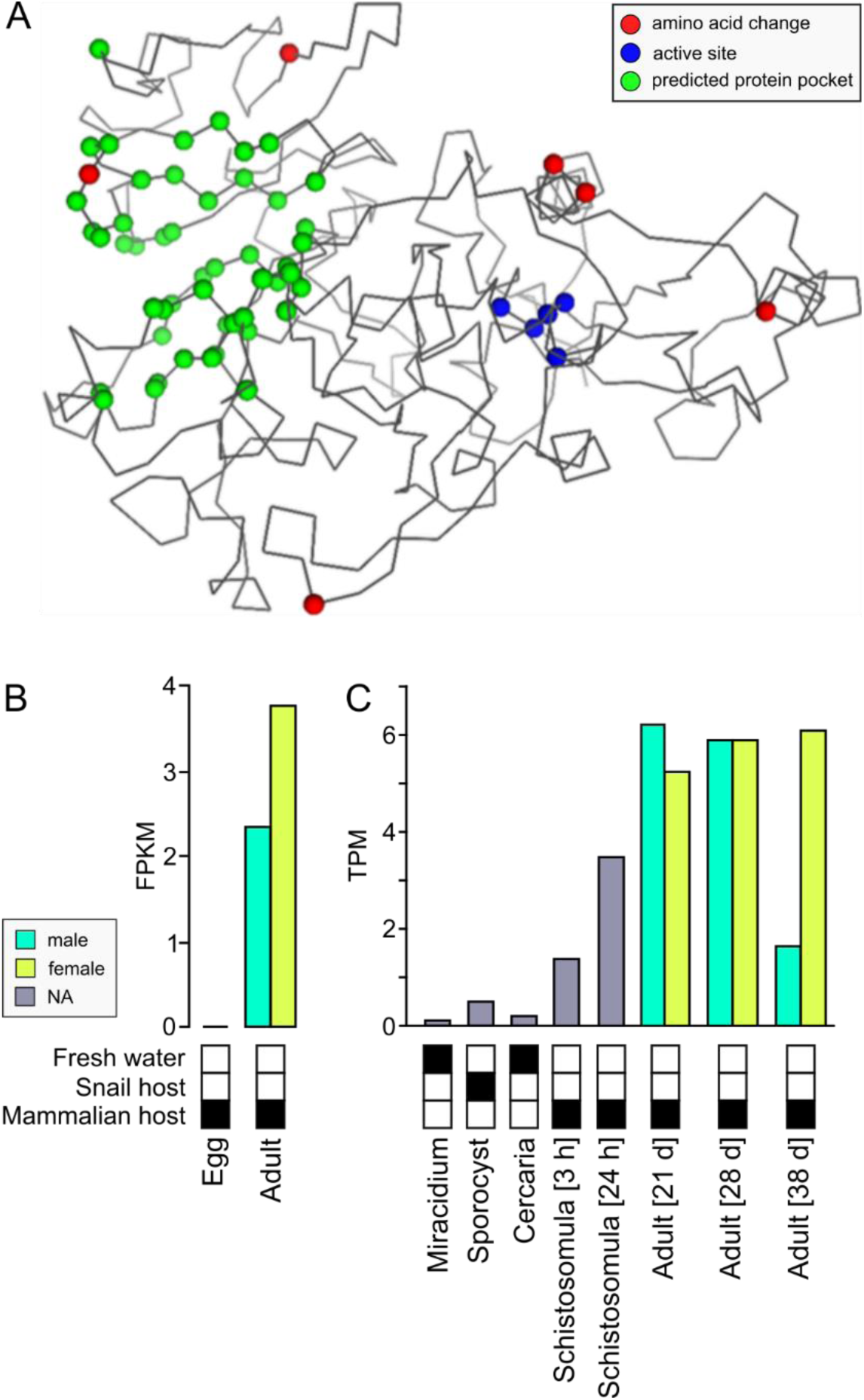
Invadolysin (Smp_127030) structure and expression. (A) The invadolysin (Smp_127030) structure was modeled based on a homologous *Drosophila* leishmanolysin structure. The amino acid changes are associated with the exterior of the protein and do not fall within the active site. (B) Invadolysin (Smp_127030) is expressed in adult worms in *S. haematobium*. (C) In *S. mansoni,* Smp_127030 is expressed in stages associated with the mammalian host, and most highly expressed in adult worms.

## Discussion

We were unable to find evidence of contemporary hybridization between *S. haematobium* and other schistosome species using samples from Zanzibar and Niger. Instead our data provide evidence for an ancient hybridization event between *S. haematobium* and *S. bovis* occurring hundred(s) of generations ago. Introgression was regional occurring in Niger but not in Zanzibar. In a Niger *S. haematobium* population 3.3-8.2% of the genome are introgressed *S. bovis* alleles. Some of the introgressed *S. bovis* alleles have almost reached fixation within *S. haematobium*, and show signatures of strong selection, providing unambiguous evidence for cross species transfer of genes through hybridization and introgression.

### Ancient introgression rather than contemporary hybridization

Multiple studies of West African *S. haematobium* miracidia have described discordant mitochondrial DNA and rDNA sequences in up to 41% of parasites sampled (Table 1). These studies have been interpreted to indicate that hybridization occurs at high frequencies within many West African locations where *S. haematobium* and *S. bovis* occur sympatrically. In contrast, our exomic analyses using samples from Niger and Zanzibar reveals ancient hybridization and subsequent introgression.

While 65% of miracidia sampled from Niger contained *S. bovis*-derived mtDNA, none of these 48 parasites showed evidence for recent hybridization based on analysis of the autosomal genome: we did not find any individual miracidia with 50% or 25% of their genome derived from *S. bovis* as would be expected for F1 or F2 hybrids. These results suggest that hybridization is either rare (0-7.4%; 95% binomial confidence interval; Clopper and Pearson 1934) in *S. haematobium*, or geographically localized. Broader geographical sampling and genomic characterization of miracidia will be needed to accurately determine the frequency of F1 or early generation hybrids in other *S. haematobium* populations.

Second, previous studies have suggested that 0-41% of parasites sampled in West African locations have hybrid origins based on mitochondrial and rDNA genotyping (Table 1). This is clearly an underestimate: we found that 100% of *S. haematobium* miracidia sampled from Niger contain *S. bovis* ancestry, with 3-8% of autosomal loci derived from *S. bovis*. We conclude that ancient hybridization and introgression best explains the Nigerien genomic data and we predict that more extensive genomic surveys will demonstrate that recent (F1 or F2) hybrids between *S. haematobium* and *S. bovis* occur rarely in nature.

Crosses between multiple members of the *S. haematobium* species group, other than *S. bovis* and *S. haematobium*, have been staged in the laboratory using rodent hosts (Webster, *et al.* 2006). Furthermore, field collected miracidia with mtDNA and rDNA markers from different species have been collected in nature. These include reports of non-viable, hybrid miracidia between *S. mansoni*, agent of intestinal schistosomiasis and *S. haematobium* (Webster, *et al.* 1999). We emphasize that our conclusions regarding ancient introgression apply to *S. haematobium* and *S. bovis* only. Genomic analyses of putative hybrid miracidia from other schistosome species combinations or locations will determine if these reflect ancient or contemporary hybridization events, and can help to understand how species barriers are maintained in schistosomes.

We estimate that the hybridization between *S. haematobium* and *S. bovis* occurred between 240 (range: 108-613) generations ago, based on the size distribution of introgressed fragments. However, we emphasize that our estimate of time since introgression is likely to be conservative because exome data provides limited ability to identify small haplotype blocks. Despite these limitations the genomic data demonstrate that the introgression event occurred more than a hundred generations ago but subsequent to the divergence of the Nigerien and Zanzibari *S. haematobium* populations. Whole genome, rather than exome data, together with improvements in the *S. haematobium* genome assembly, and broader geographical sampling will improve our dating estimates of ancient hybridization with *S. bovis*.

Given that exome capture probes failed to capture *S. bovis* mtDNA, there is also a potential concern that our exome capture approach will preferentially capture *S. haematobium* DNA, and underestimate *S. bovis* alleles within the nuclear genome. We do not think this is the case for two reasons. First, *S. haematobium* and *S. bovis* genomes are ∼3% divergent (Oey, *et al.* 2019). Exome capture effectively captures genomes at this level of divergence with minimal bias (Bi, *et al.* 2012). Second, we examined proportions of *S. bovis* ancestry in 12 miracidia for which collected both exome and genome sequence data (Fig S4). Ancestry estimates were consistent across both exome and genome sequence data with the largest difference between estimates being 0.015 (1.5%).

### Introgression is geographically widespread

How widespread is this ancient introgression event in Africa? Our results indicate regional introgression in parasites from Niger, but not from Zanzibar. Introgression may extend across multiple countries in Africa. Oey *et al*. (2019) sequenced the *S. bovis* genome and compared this with the Egyptian *S. haematobium* reference. They noted high similarity between *S. bovis* and the *S. haematobium* reference strain in genome regions spanning up to 100Kb and hypothesized that these regions were the result of an introgression event between *S. bovis* and *S. haematobium*. Interestingly, one of these high similarity segments is on chromosome 4 and spans the invadolysin locus (Smp_127030). Taken together, our findings, in combination with Oey, *et al.* (2019), indicate that the common ancestor of the Nigerien and Egyptian *S. haematobium* reference hybridized with *S. bovis,* suggesting that remnants of this introgression event may be widespread in extant *S. haematobium* in Africa. Interestingly, *S. bovis* has been recorded in Zanzibar (Pennance, *et al.* 2018), but we see no evidence for introgression of *S. bovis* alleles in Zanzibari *S. haematobium*. Further, sampling will reveal the number and geographical extent of introgression events in *S. haematobium* across Africa.

### The nature of selection driving introgression

Invadolysin is a member of a M8 metalloprotease gene family originally identified in *Leishmania* (Russell and Wilhelm 1986). In *Leishmania*, these are surface proteins associated with degradation of collagen and fibrenonectin in the extracellular matrix and larval penetration of mammalian hosts (Yao, *et al.* 2003). In schistosomes M8 metalloproteases have undergone a rapid gene family expansion (Figure S3) and are among the most abundant transcripts and secreted proteins (Curwen, *et al.* 2006; Liu, *et al.* 2014; Coghlan, *et al.* 2019) in larval stages. RNAi knockdowns of one invadolysin paralog in miracidia of *S. mansoni* led to reduced larval penetration and establishment in the snail intermediate host, reducing cercariae production (Hambrook, *et al.* 2018).

Members of the invadolysin family are expressed in a stage specific manner, similar to globins in vertebrates (Storz, *et al.* 2013). We suspect that the introgressed *S. bovis-*derived invadolysin is selected during the mammalian, rather than the snail infective stage of the lifecycle. The *S. mansoni* paralog of this gene (Smp_127030) is expressed during stages associated with the mammalian host and most highly expressed in adult worms (Figure S5). Consistent with this, the Smp_127030 is expressed in adult worms in *S. haematobium* (Figure 7B), but expression data is not available for other life stages (except eggs). It is unclear how the *S. bovis* invadolysin (Smp_127030) allele benefits *S. haematobium* based on the currently available data. Our exome probes covered half of the invadolysin (Smp_127030) exons, and fewer than half of the non-synonymous SNPs we identified were able to be placed on the inferred protein structure. Despite these limitations, we find strong evidence of selection from the available population data. We speculate that the introgressed invadolysin is involved in tissue penetration (Wilson 2012) or immune evasion (Hambrook, *et al.* 2018) within the mammalian host. Functional analysis of this system may be possible, since certain strains of *S. haematobium* can be maintained in the laboratory, and will be a central priority for future work with this system.

### Implications for schistosome control

Significant resources are being directed towards schistosome control and elimination strategies (Savioli, *et al.* 1997; Zhou, *et al.* 2005; King 2009; Webster, *et al.* 2014; Webster, *et al.* 2016; Tchuenté, *et al.* 2017). Hybridization between human and livestock schistosomes, including *S. bovis* and *curassoni*, will complicate these efforts (King, *et al.* 2015). The introgression of the *S. bovis* invadolysin allele in *S. haematobium* provides an example of an adaptive introgression between an animal and human parasite species. Gene exchange between these two species highlights the need for effective coordination between medical and veterinary researchers and a “one health” approach to schistosome control and management (Webster, *et al.* 2016).

## Conclusions

Hybridization between species is a powerful source of evolutionary novelty. In humans up to 4% and 5% of the genome contains Neanderthal (Green, *et al.* 2010) or Denisovan (Reich, *et al.* 2010) ancestry as a direct result of admixture with these archaic species. Genes involved in immune response (Abi-Rached, *et al.* 2011), pathogen defense (Enard and Petrov 2018), and protection from sun exposure (Dannemann and Kelso 2017) have introgressed from archaic humans as a result of these ancient hybridization events. One of the most striking conclusions from the many studies on human/Neanderthal or human/Denisovan admixture is that limited introgression can have significant genomic and phenotypic impacts.

Gene transfer across species boundaries has clearly taken place in *S. haematobium*. While we see no evidence of early generation hybrids, we see a signature of ancient introgression from *S. bovis* in all Nigerien miracidia examined. We demonstrate that an introgressed M8 metalloprotease (invadolysin; Smp_127030) expressed in adult worms has spread to high frequencies in *S. haematobium* populations in Niger, and that parasites bearing introgressed *S. bovis*-derived alleles have a strong selective advantage over those carrying native *S. haematobium* alleles. Understanding the parasite phenotype conferred by the introgressed *S. bovis* alleles and the nature of selection driving the spread of these *S. bovis* alleles into *S. haematobium* populations is a high priority.

## Materials and Methods

### Ethics statement

For the Niger sample collection, ethical clearance was obtained from the Niger National Ethical Committee, in combination with the St Mary’s Hospital Local Ethics Research Committee (part of the Imperial College London Research Ethics Committee (ICREC; (EC NO: 03.36. R&D No: 03/SB/033E)) in London, United Kingdom, in combination with the ongoing Schistosomiasis Control Initiative (SCI) and Schistosomiasis Consortium for Operational Resaerch (SCORE) activities. For the Zanzibar sample collection, ethical approval was obtained from the Zanzibar Medical Research and Ethics Committee (ZAMREC, reference no. ZAMREC 0003/Sept/011) in Zanzibar, United Republic of Tanzania, the “Ethikkomission beider Basel” (EKBB, reference no. 236/11) in Basel, Switzerland, the Institutional Review Board of the University of Georgia (project no. 2012-10138-0), and registered at the International Standard Randomised Controlled Trial (Register Number ISRCTN48837681). Within both Niger and Zanzibar, all aspects of sample collections were carried out in the framework of the disease control activities implemented and approved by the local Ministry of Health and adopted by regional and local administrative and health authorities.

The study participants were informed about the study objectives and procedures. Written consent was obtained from parents prior to sample collection from children. Participation was voluntary and children could withdraw or be withdrawn from the study at any time without obligation. All children were offered PZQ (40 mg/kg single oral dose) treatment in the frame of the following school-based or community-wide treatment carried out by the Ministry of Health.

### Sample collection

This study used archived miracidia samples from Niger and Zanzibar (Tanzania) fixed on Whatman FTA cards archived within the <u>S</u>chistosome <u>C</u>ollection <u>a</u>t the <u>N</u>atural History Museum (SCAN)(Emery, *et al.* 2012). From Zanzibar, encompassing both the Unguja and Pemba islands, the *S. haematobium* miracidia were collected as part of the SCORE population genetics studies in 2011 from 26 locations spaced up to 160.9 km apart. From Niger the *S. haematobium* miracidia were collected in 2013 from school-aged children from 10 locations located up to 125 km apart. These samples were also collected as part of SCORE population genetic studies within a gaining and sustaining control of schistosomiasis project in Niger and also as part of monitoring and evaluation activities carried out by the Schistosomiasis Control Initiative. To capture maximum diversity we used a single miracidium from 96 individuals (*n*_Zanzibar_=48, *n*_Niger_ = 48). *S. haematobium* eggs were harvested from each infected urine sample by sedimentation or filtration, put into fresh water and then exposed to light to facilitate miracidial hatching. Miracidia were captured individually, under a binocular microscope, in 3 µL of water and spotted onto indicating Whatman FTA Classic Indicating cards (GE Healthcare Life Sciences, UK) using a micropipette, dried and archived in SCAN(Emery, *et al.* 2012).

### Library prep and sequencing

We used whole genome amplification of single miracidia dried on FTA cards followed by exome capture and sequencing (Illumina 2500) to generate genome-wide sequence data following methods described in Le Clec’h et al ((Le Clec’h, *et al.* 2018)). The exome capture probe set (SureSelect design ID: S0742423) was designed using the published reference genome sequence for *S. haematobium* (SchHae_1.0, GCA_000699445.1, lab strain, originally from Egypt). The exome capture probe set included 156,004,120bp RNA baits from the nuclear genome and 67 from the mitochondrial genome, covering 96% (62,106/64,642) of the exons and accounting for 94% of the exome length (15,002,706 bp/15,895,612 bp). Each captured exon was covered by an average of 2.59 120bp baits (Le Clec’h, *et al.* 2018).

In addition to exome capture, we generated whole-genome sequence data for twelve samples, six from each population (Niger and Zanzibar). Libraries were multiplexed and paired end 150 bp reads were sequenced on a single Illumina NextSeq flowcell. In addition to the whole genome and exome sequence data generated above, we gathered available genome sequence data from the NCBI Short Read Archives for six other species of *Schistosoma* in the *haematobium* group, including *S. bovis* (ERR119622, ERR103048, ERR539853), *S.* c*urassoni* (ERR310937, ERR119623), *S. haematobium* (ERR084970, ERR037800, SRR433865), *S. intercalatum* (ERR539854, ERR539856, ERR119613), *S. guineensis* (ERR119612, ERR539850, ERR539852), *S. mattheei* (ERR539851, ERR539855, ERR539857, ERR103051), and *S. margrebowiei* (ERR310940) in 20 separate libraries(Young, *et al.* 2012; Coghlan, *et al.* 2019).

### Computational environment

We conducted all analyses on a high-performance computing cluster within a Singularity container or Conda environment. Environmental recipe files, custom programs, and shell scripts are at https://github.com/nealplatt/sH_hybridization (v1.0; doi: 10.5281/zenodo.2536390).

### Variant discovery, filtration, and phasing

We trimmed sequence reads with Trimmomatic v0.36 (Bolger, *et al.* 2014) so that the leading and trailing base calls had phred-scaled quality scores greater than 10 and the phred score was greater than 15 over a 4 nt sliding window. After trimming, we mapped paired and singleton reads to the reference *S. haematobium* genome (SchHae_1.0, GCA_000699445.1, lab strain, originally from Egypt). BWA v0.7.17-r1188 (Li and Durbin 2009) and merged into a single BAM file using SAMtools v1.7 (Li, *et al.* 2009). GATK v4.0.1.1 (McKenna, *et al.* 2010) was used to add read group information and mark duplicate reads from each library. Where possible, we added complete read group information based on information contained within the FASTQ header. In some cases (primarily the public data) pseudo read groups were created to differentiate each library.

We used GATK’s Haplotypecaller (McKenna, *et al.* 2010) for variant discovery. Variant discovery was restricted to target regions of the *S. haematobium* assembly using the –L option. Target regions were identified by combining all adjacent, exome probe locations within 500 bases of one another into larger intervals with BEDtools v2.27.1 (Quinlan and Hall 2010). Each interval was genotyped using GATK’s GenotypeGVCFs and combined into a single VCF for filtering using GATK’s MergeVcfs. Low quality SNP genotypes were filtered using GATK’s VariantFiltration with the following filters: variant quality normalized by depth (QD < 2.0), strand bias (FS > 60.0), mapping quality (MQ < 40.0), mapping quality rank sum (MQRankSum < −12.5), and read position rank sum (ReadPosRankSum < −8.0). We used VCFtools v 0.1.15(Danecek, *et al.* 2011) to remove sites with high rates of missing data (>20%), multi-allelic sites, and individuals with low call rates (>15% missing data). All indels were excluded from downstream analyses.

The published *S. haematobium* and *S. mansoni* assemblies vary greatly in quality, despite a high degree of synteny between the two genomes (Young, *et al.* 2012). We aligned the two genomes using progressiveCactus v0.0 (Paten, *et al.* 2011) using default parameters to leverage the contiguity of the *S. mansoni* assembly. The HAL alignment file was used to lift SNP coordinates between the assemblies using progressiveCactus’ halLiftover module. We removed multi-position SNPs, those that align between assemblies in something other than a 1:1 relationship, from downstream analyses. We used linkage disequilibrium (LD) decay curves to examine biases introduced during the coordinate liftover by comparing LD from SNPs associated with each assembly. SNPs mapping to the *S. mansoni* autosomes (chr1-7) were extracted using VCFtools v0.1.15. The square of the correlation coefficient (*r^2^*) was calculated for all SNPs within 1.5 Mb of each other for each dataset using PLINK v1.90b4 (Purcell, *et al.* 2007). The 1.5 Mb cutoff was choosen since it represented the largest scaffold in the *S. haematobium* assembly. After confirming concordance between the original and *S. mansoni*-lifted datasets, we used Beagle v4.1 (Browning and Browning 2016) to impute missing SNPs and phase haplotypes for each autosomal chromosome. Imputations and phasing occurred in sliding windows of 300 SNPs and a step size of 30 with 250 iterations per window.

### Mitotype assignment

Unique filters were used to manage the mitochondrial SNPs. First, the mitochondrial contig was identified from the *S. haematobium* assembly using the *haematobium* vs. *mansoni* whole genome alignment and confirmed using NCBI’s BLAST server. Due to low genotyping rates within the Nigerien samples, mitochondrial SNPs were filtered so that the genotyping rate per site was reduced from 85% to 25%. Putative mitochondrial haplotypes (mitotypes) were generated by removing any heterozygous sites, if present. Given previously described rates of mitochondrial introgression and divergence between *S. bovis*, *S. curassoni* and *S. haematobium,* we mapped filtered reads directly to the *S. haematobium* mitochondrial contig (AMPZ01026399.1) and a previously generated *S. curassoni* mitochondria sequence (AP017708.1). Full-length *S. bovis* mitochondrial genomes are not currently available through NCBI Genbank (last accessed 23 April 2018). *S. bovis*/*curassoni* or *haematobium* mitotypes were classified using the ratio of reads mapping to each mitochondrial reference and confirmed by Sanger sequencing the mitochondrial, cytochrome C oxidase 1 (*cox1*) gene. *cox1* was amplified in a reaction containing 1x reaction buffer, 0.8 mM dNTPs, 1.5mM MgCl2, 1 µM forward primer (Cox1_schist_5’; CTT TRG ATC ATA AGC G), and 1 µM reverse primer (Cox1_schist_3’; TAA TGC ATM GGA AAA AAA CA) and under the following reaction conditions: 2 minute (min), 95°C initial denaturation, 35 cycles of a 30 second (sec), 95°C denaturation, 30 sec, 52°C annealing phase, and a 1 min 72°C extension, followed by a final 7 min, 72°C extension cycle (Lockyer, *et al.* 2003). Amplification and fragment size were confirmed on a 1.5% agarose gel. Bidirectional sequences were generated for each fragment with the Cox1_schist_5’ and Cox1_schist_3’ primers by the Eurofins (Eurofins-MWG) sequencing service. All *cox1* sequences were compared to available sequences on GenBank for species identification.

### Population structure, admixture, and ancestry assignment

Summary statistics were calculated using all filtered autosomal SNPs with minor allele frequency greater than 5%. FIS was calculated for each sample using VCFtools and the “--het” option. The scikit-allel v1.1.10 (Consortium 2017) Python library was used to calculate F_ST_ between the Zanzibari and Nigerien populations of *S. haematobium*. Genome-wide values for F_ST_ were averaged from blocks of 100 variants and local calculations were generated from sliding windows of 250 kb and 50 kb steps. Relationships between samples were visualized in genotypic space with a PCA of unlinked SNPs. Linked sites within sliding windows of 25 SNPs and a pairwise R^2^ value greater than 0.2 were filtered using PLINK v1.90b4 (Purcell, *et al.* 2007). Since comparison between species can overwhelm population level clustering, separate PCAs were generated from previously published *Schistosoma* species (Young, *et al.* 2012; Coghlan, *et al.* 2019), and the exome data from the *S. haematobium* miracidia (Niger and Zanzibar) presented herein. Nucleotide diversity (π) was calculated per site and in sliding windows of 50 Kb with 25 Kb steps using VCFtools.

Levels of admixture were calculated using ADMIXTURE v1.3.0 (Alexander, *et al.* 2009). The number of populations (*K*) was explored from 1 to 20 and cross-validation between 1,000 replicates was used to identify the most robust measure of *K*. The 3-population test *f_3_* as implemented in the scikit-allel was used to identify admixture in the Nigerien and Zanzibar *S. haematobium* populations (Reich, *et al.* 2009). *F_3_*, standard errors, and *Z*-scores were calculated from block-jackknife replicates of 100 SNPs. We used the scikit-allel package to calculate *D* statistics and compare possible introgression from *S. bovis* or S. *curassoni* into *S. haematobium*. Standard error and *Z*-scores were calculated by block-jackknife replicates in windows of 100 SNPs. In addition to genome-wide averaged, *D* was calculated in local, sliding windows of 50 SNPs with 25 SNP steps. Finally, PCAdmix v1.0 (Brisbin, *et al.* 2012) was used for SNP ancestry assignment. For individuals in the admixed Nigerien population, ancestry was assigned in sliding windows across each chromosome to one of two parental populations defined by “pure” *S. haematobium* (represented here by samples from Zanzibar) and *S. bovis*. For validation, we included five, randomly selected, Zanzibari *S. haematobium* samples with the assumption that the entire genome should be assigned Zanzibari ancestry. Ancestry was assigned to windows of 30 SNPs at a time, with a minimum assignment threshold of 99%.

### Selection

We quantified selection across the *S*. *haematobium* exome using two independent methods. BayeScan v2.1 (Foll and Gaggiotti 2008) proposes two demographic models to explain allele frequency differences between at each locus. The first model assumes allele frequency differences are due to drift while the other model adds an additional parameter (α) to account for selection. A Markov-chain Monte Carlo (MCMC) was used to generate a posterior distribution of the alpha parameter such that significant deviations from zero indicate directional selection. We examined selection by comparing the Zanzibar and Niger *S. haematobium* populations using 1,000 separate BayeScan runs (chains) to guarantee convergence on posterior distributions alpha. For each chain we used 50 pilot runs of 10,000 generations to generate starting priors and burnin time of 50,000 generations. Prior odds for the neutral models (α = 0) were set at 10:1. To minimize autocorrelation, samples were thinned by 20 and 50,000 samples were taken. Convergence between chains was examined in R using the *coda* package v0.19-1. A Gelman-Rubin convergence diagnostic score less than 1.1 was defined as an acceptable level of convergence *a priori*. In addition to BayeScan, we identified regions under selection in both the *S. haematobium* populations using cross population extended haplotype homozygosity ((xpEHH; Sabeti, *et al.* 2007)) score as implemented in the *rehh* v2.0.2 R package (Gautier and Vitalis 2012). When possible, alleles were polarized against an out-group (*S. margrebowiei*) to define the ancestral state. If sites were heterozygous or missing in *S. margrebowiei* they were excluded. Regions under selection were defined using a -log_10_ (*p*-value) = 3 an an |xpEHH| > 3. Adjacent loci ≤12.5 kb apart were combined into a single locus.

### Dating genome wide introgression

*S. bovis* introgression tracts were identified using Python scripts available at https://github.com/nealplatt/sH_hybridization (release v1.0; doi: 10.5281/zenodo.2536390). Briefly, derived (autapomorphic) *S. bovis* and Zanzibari *S. haematobium* alleles were identified and used to annotate Nigerien *S. haematobium* alleles. Introgression blocks were identified by finding the largest stretch of *S. bovis* or pleisiomorphic alleles that were unbroken by a Zanzibari *S. haematobium* allele. With these tracts we estimated the number of generations since admixture (Schumer, *et al.* 2016):

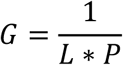

Where *G* is generations, *L* is the average length of introgression tracts in Morgans and *P* is the proportion of the genome from the major parent.

We dated time since divergence of the Chr4:20,023,951-20,047,325 locus in the Nigerien *S. haematobium* populations using startmrca (Smith, *et al.* 2018). We used a uniform recombination rate of 3.4 × 10^−8^ and mutation rate of 8.1 × 10^−9^. We centered the analysis on Chr4:20,033,013 and included 1Mb of upstream and downstream sequence. MCMC chains were run for 50,000 generations and limiting proposals to 20 standard deviations. Ten independent chains were run using the parameters described above. Estimates of divergence time were taken by discarding the first 40,000 generations of each run then thinning the remainder to 1 sample per 10 generations.

### M8 peptidase and Invadolysin gene family evolution

Invadolysin (Smp_127030) is a member of the M8 peptidase family of proteins (Pfam ID: PF01457) from the Pfam database (v32.0; last accessed 28 October 2018; Finn, *et al.* 2013). We downloaded amino acid sequences for all platyhelminths and several outgroup taxa including *Homo sapiens*, *Mus musculus*, *Monodelphis Domestica*, *Gallus gallus*, *Anolis carolinesis*, *Danio rerio*, *Drosophila melanogaster*, *Aedes* aegypti, and *Caenorhabditis elegans*. Sequences missing the HExxH active site (Rawlings and Barrett 1995) were removed from downstream analysis. Amino acid sequences were aligned with MUSCLE v3.8.1551 (Edgar 2004), and sites with less than 75% coverage were removed. We used RAxML v 8.2.10 (Stamatakis 2014) to complete 100 searches for the optimal tree using the PROTGAMMAWAG substitution model. Gene duplication evens were inferred using Mega7 (Kumar, *et al.* 2016) and the gene duplication wizard. Duplications were categorized based on their presence in lineages leading to *Schistosoma* paralogs.

### Invadolysin (Smp_127030) structure

We used Phyre2 (Kelley, *et al.* 2015) in “intensive mode” to model the Nigerien *S. haematobium* invadolysin (Smp_127030) protein structure. We used the whole genome resequencing data to identify SNPs at the invadolysin (Smp_127030) locus since the exome probes only targeted 76.1% of the coding region (1,722 of 2,262 bp). Pockets in the protein structure were identified with fpocket2 (Le Guilloux, *et al.* 2009) as implemented in PHYRE2 web portal. After modeling the invadolysin (Smp_127030) protein structure, we examined the placement of amino acid differences fixed between the Zanzibari and Nigerien *S. haematobium* populations with PyMol v2.2.3 (DeLano 2002).

### Invadolysin (Smp_127030) expression

We quantified gene expression of Smp_127030 in *S. haematobium* using previously published data (Young, *et al.* 2012) from schistosome eggs, and adult worms from both sexes (SRA accessions: SRX3632877, SRX3632879, SRX3632881). Reads were mapped to the *S. haematobium* genome (SchHae_1.0) using hisat v2.1.0 (Kim, *et al.* 2015). Transcripts were assembled and gene expression was normalized and quantified using Stringtie v1.3.4 (Pertea, *et al.* 2015). Gene coordinates were lifted from the *S. mansoni* assembly to identify which of the Stringtie predicted transcripts were invadolysin (Smp_127030) using progressiveCactus’ halLiftover and the whole genome alignment generated (described above).

Given the limited gene expression data available for *S. haematobium*, we examined expression of Smp_127030 and other invadolysin paralogs in *S. mansoni* (Berriman, *et al.* 2009). We used RNA-seq data from miracidia (Wang, *et al.* 2013), cercariae (Protasio, *et al.* 2012), schistosomula (3h, 24h, in vitro transformation), sporocysts (48h in vitro transformation), juveniles (single sex; 18, 21 28 days old), and adults (single sex, mixed infections; 38 days). (Protasio, *et al.* 2012; Wang, *et al.* 2013; Protasio, *et al.* 2017). We aligned the data using STAR v2.5.4b(Dobin, *et al.* 2013). STAR references were prepared using the v7 *Schistosoma mansoni* genome the related v7.1 annotation (Coghlan, *et al.* 2019) which was converted to the GTF format using gffread from cufflinks v2.2.1 (Trapnell, *et al.* 2010) and either an overhang (-sjdbOverhang) of 75 (used for schistosomula and cercariae data) or 99 (used for all other libraries). We used RSEM (Li and Dewey 2011) to compute transcript per million (TPM) counts for each isoform.

## Data Availability

Sequence data are deposited in NCBI’s under BioProject accessions PRJNA508633 and PRJNA443709. Mitochondrial genomes are available in are deposited in Genbank (MK253567-MK253578).

## Code Availability

Scripts, code, and environmental files are available at https://github.com/nealplatt/sH_hybridization (release v1.0; doi: 10.5281/zenodo.2536390).

## Acknowledgements

Sandy Smith, Richard Polich and Roy Garcia (Texas Biomedical Research Institute) provided research support. Steffi Knopp (Swiss Tropical and Public Health Institute) Khalfan Mohammed, Mtumweni Ali Muhsin (Helminth Control Laboratory, Unguja) and Mr. Haji Khatib Faki (Public Health Laboratory, Pemba) for their assistance in the collection of the Zanzibar samples. Matthew Berriman and collegues provided genome sequence data prior to publication. Muriel Rabone for mantiaining the SCAN database. This research was funded by the National Institute of Allergy and Infectious Diseases (R01 AI097576-01; 5R21AI096277-01), the National Center for Advancing Translational Sciences (UL1TR001120), Texas Biomedical Research Institute Forum (award 0467), the National Center for Research Resources (C06 RR013556; RR017515), the Wellcome Trust (104958/Z/14/Z), the Gates Foundation coordinated by the University of Georgia Research Foundation Inc (RR374-053/5054146; RR374-053/4785426), and a ZELS research grant (combined BBSRC, MRC, ESRC, NERC, DSTL and DFID: BB/L018985/1). WL was supported by a Cowles Fellowship.

## Author contributions

Study Design (T.J.C.A., J.P.W, D.R., A.M.E., B.L.W.), Formal Analysis (R.N.P.II, T.J.C.A., W.L., F.D.C), writing – original draft (R.N.P.II, T.J.C.A); writing – review and editing (M.M.W, W.L, F.D.C, B.L.W, J.P.W, D.R, A.M.E); sampling (J.P.W, D.R., F.A., B.L.W, A.G, F.K, S.M.A, A.M.E).

---

**Supplemental Figure S1.**
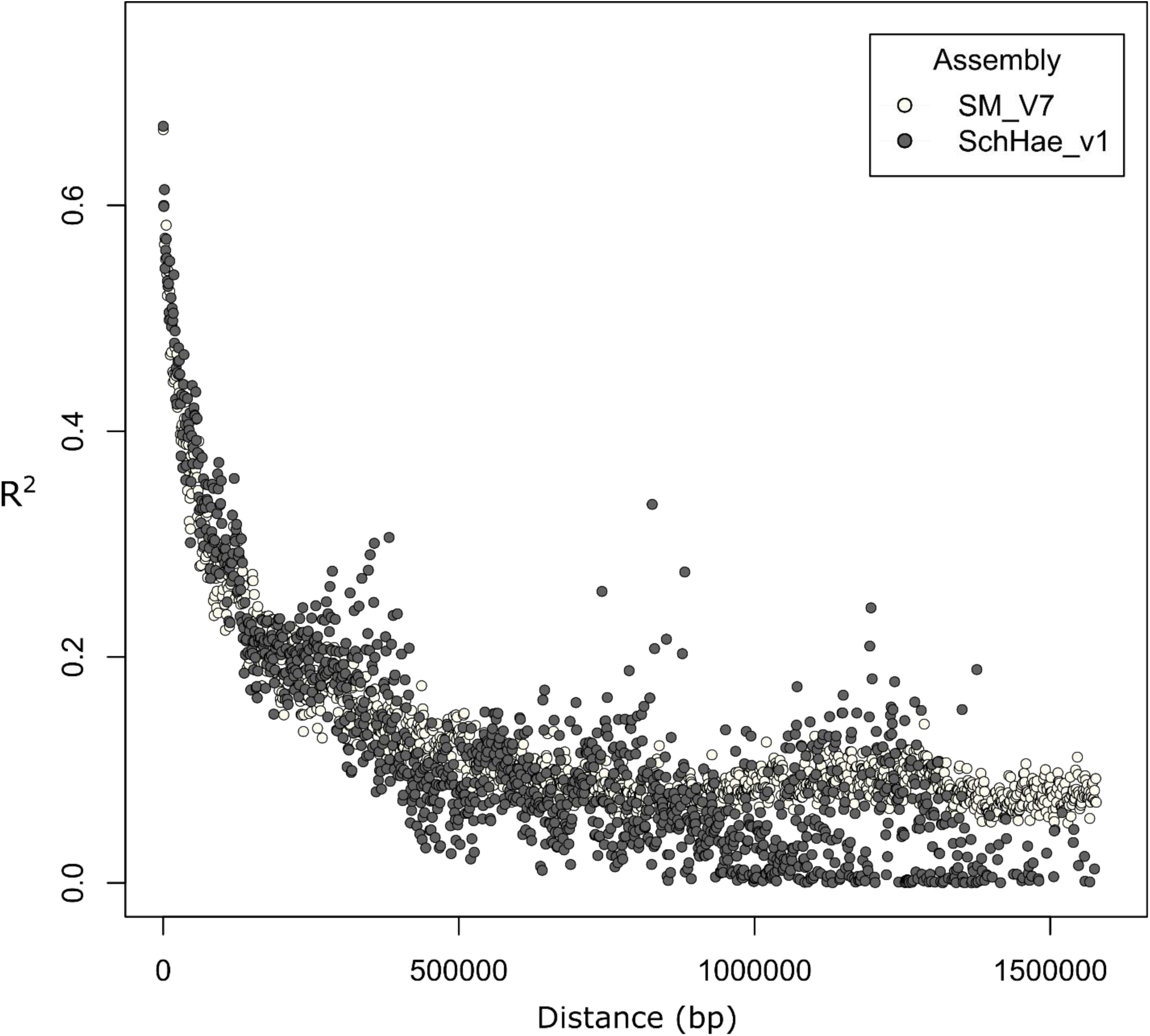
Comparison of linkage disequilibrium (LD) between different genome assemblies. SNPs were genotyped using the current *Schistosoma haematobium* assembly (SchHae_v1). The *S. mansoni* genome (SM_V7) is a more contiguous assembly with most sequences contained in chromosomal length scaffolds. To take advantage of this contiguity, SNP coordinates were lifted from *S. haematobium* to the *S. mansoni* coordinates. LD was quantified from the *S. haematobium* SNPs on both assemblies. LD decays at similar rates for each set of coordinates, indicating a degree of synteny between the two genomes.

**Supplemental Figure S2.**
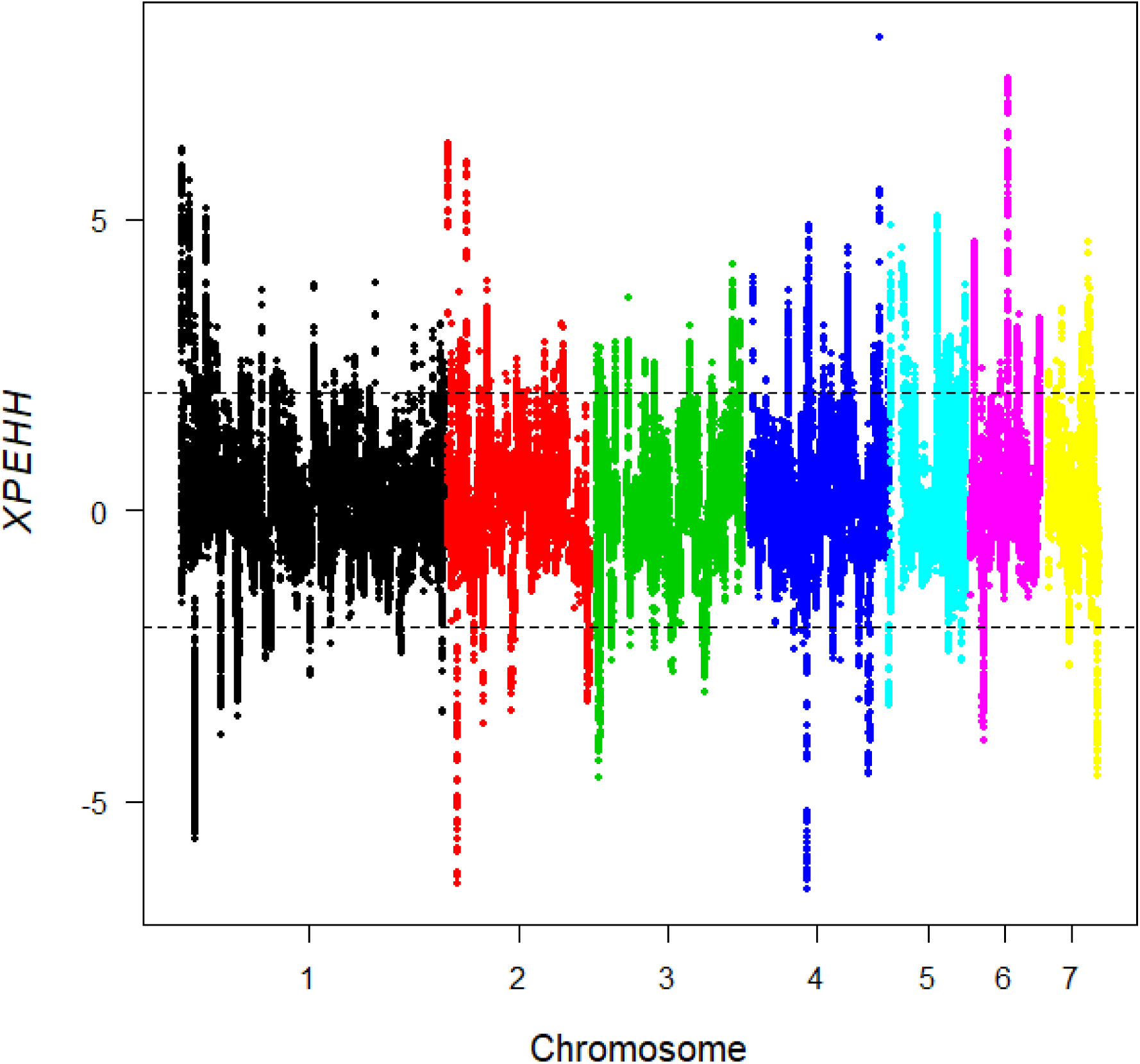
Selection across the genome between Zanzibari and Nigerien *S. haematobium* populations. Selection across the genome was measured using cross population extended haplotype homozygosity (xpEHH). xpEHH > 2 indicates directional selection in the Zanzibari population and xpEHH < −2 indicates directional selection in the Nigerien *S. haematobium* population. Strong signals of selection are present throughout the genome, but the strongest signal of directional selection in the Nigerien population is on Chr4 and spans the invadolysin (Smp_127030) gene.

**Supplemental Figure S3.**
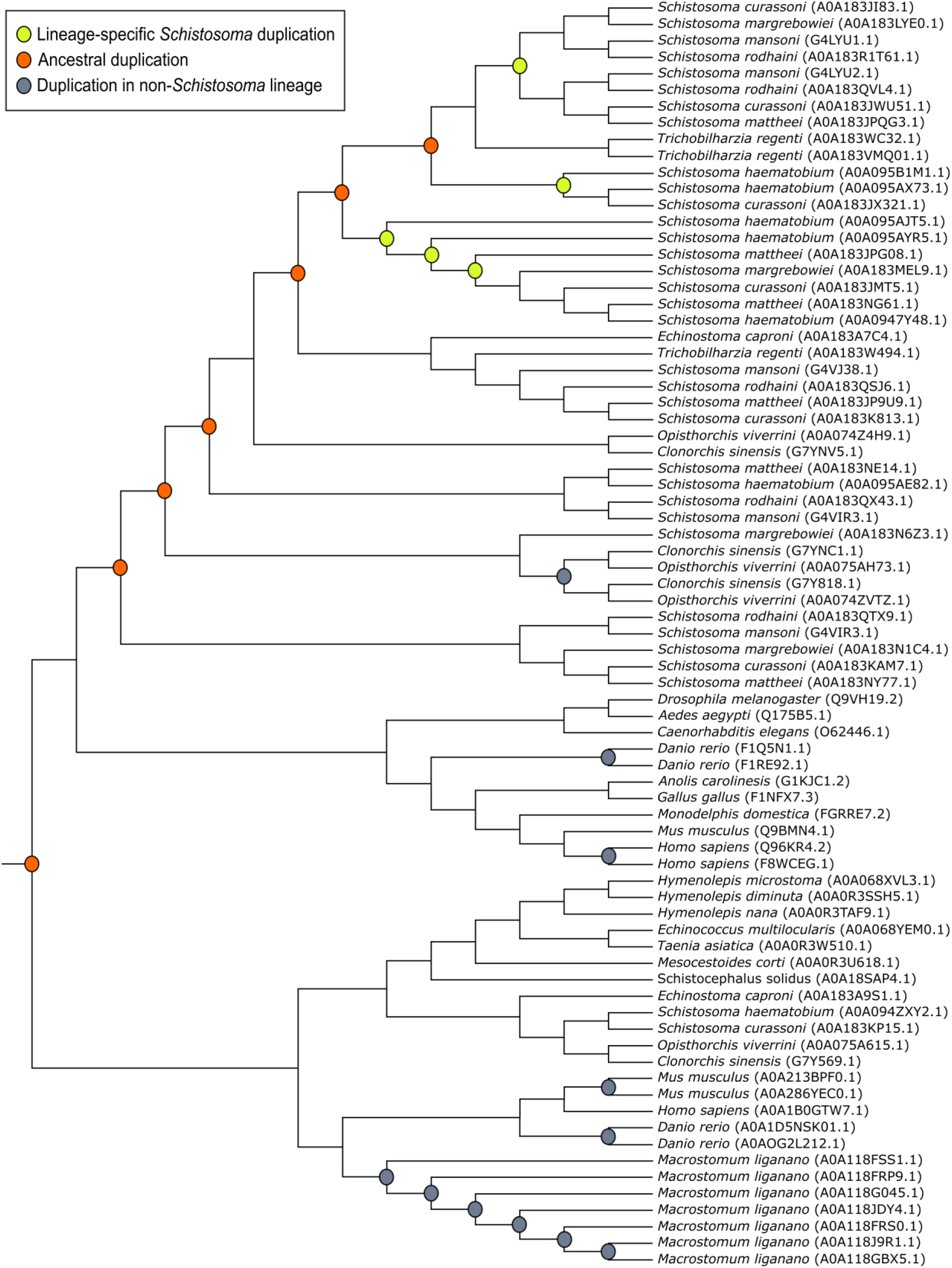
A phylogenetic tree of M8 metalloproteases, including invadolysin (Smp_127030), in platyhelminths and selected outgroup species. Gene duplications are shown at each node with a circle and colored based on its presence in Schistosoma. Of the 77 sequences examined, paralogs in *S*. *curassoni*, *haematobium*, *mansoni*, *margrebowiei*, *mattheei*, and *rodhaini* accounted for almost half (n = 34). Duplications were classified based on the presence of schistosomes at the termini. Ancestral duplications are at nodes containing schistosomes and other taxa, lineage specific duplications are at nodes containing only schistosomes, and duplications at nodes without schistosomes. Multiple, lineage-specific gene duplications are necessary to explain the distribution of metalloproteases in *Schistosoma*.

**Supplemental Figure S4.**
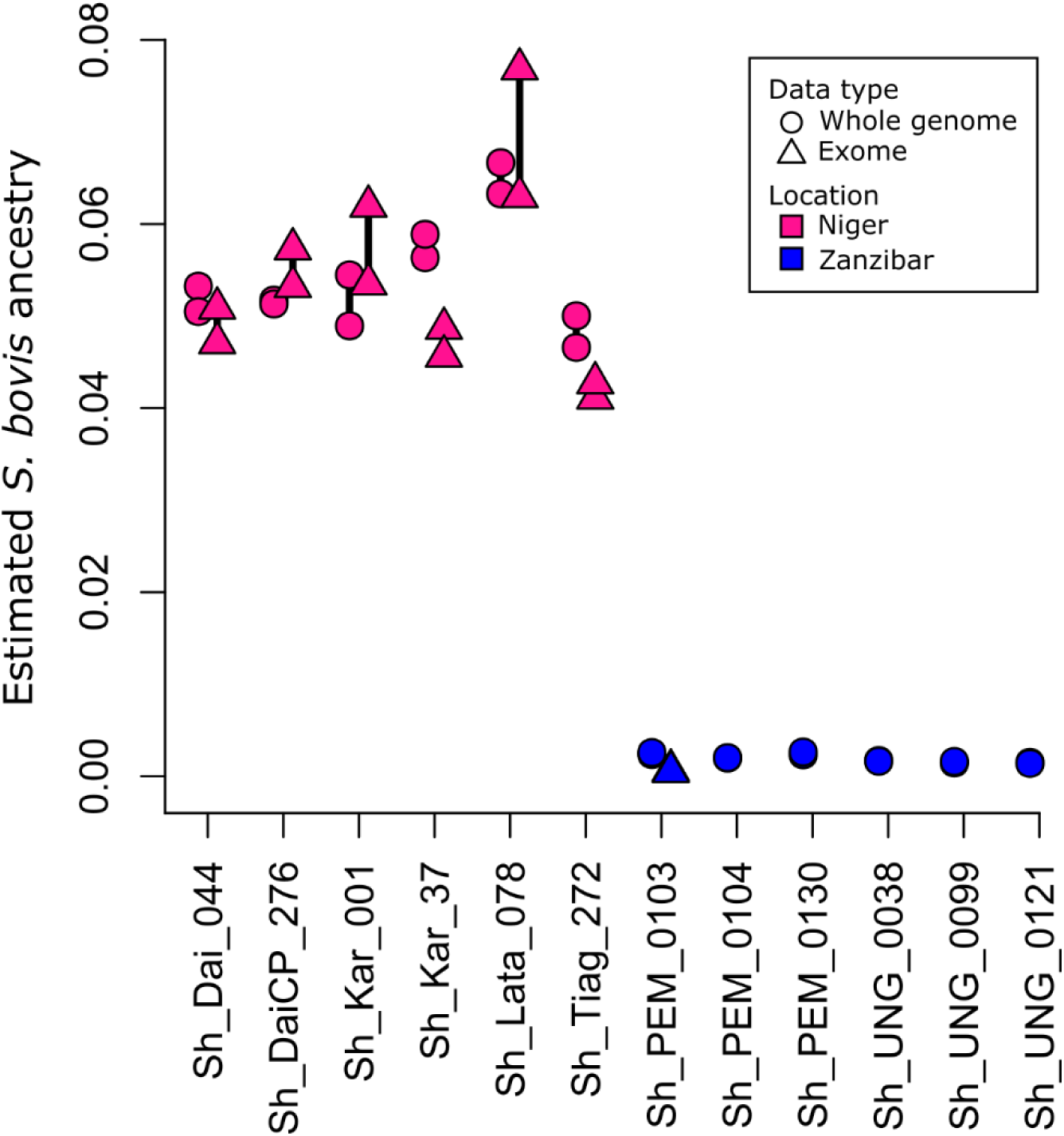
Comparing *S. bovis* ancestry estimates between whole genome and exome data. Whole genome and exome data was generated for 12 *S. haematobium* individuals. To identify potential biases from either data type we compared the *S. bovis* ancestry estimates generated by PCAdmix. Ancestry estimates from exome data were available for all Nigerien and one Zanzibari *S. haematobium*. Ancestry estimates from exome data were not available for five Zanzibari *S. haematobium* since they were designated in the parental population. Ancestry estimates were consistent across both data types with the largest variation being 0.015 (1.5%).

**Supplemental Figure S5.**
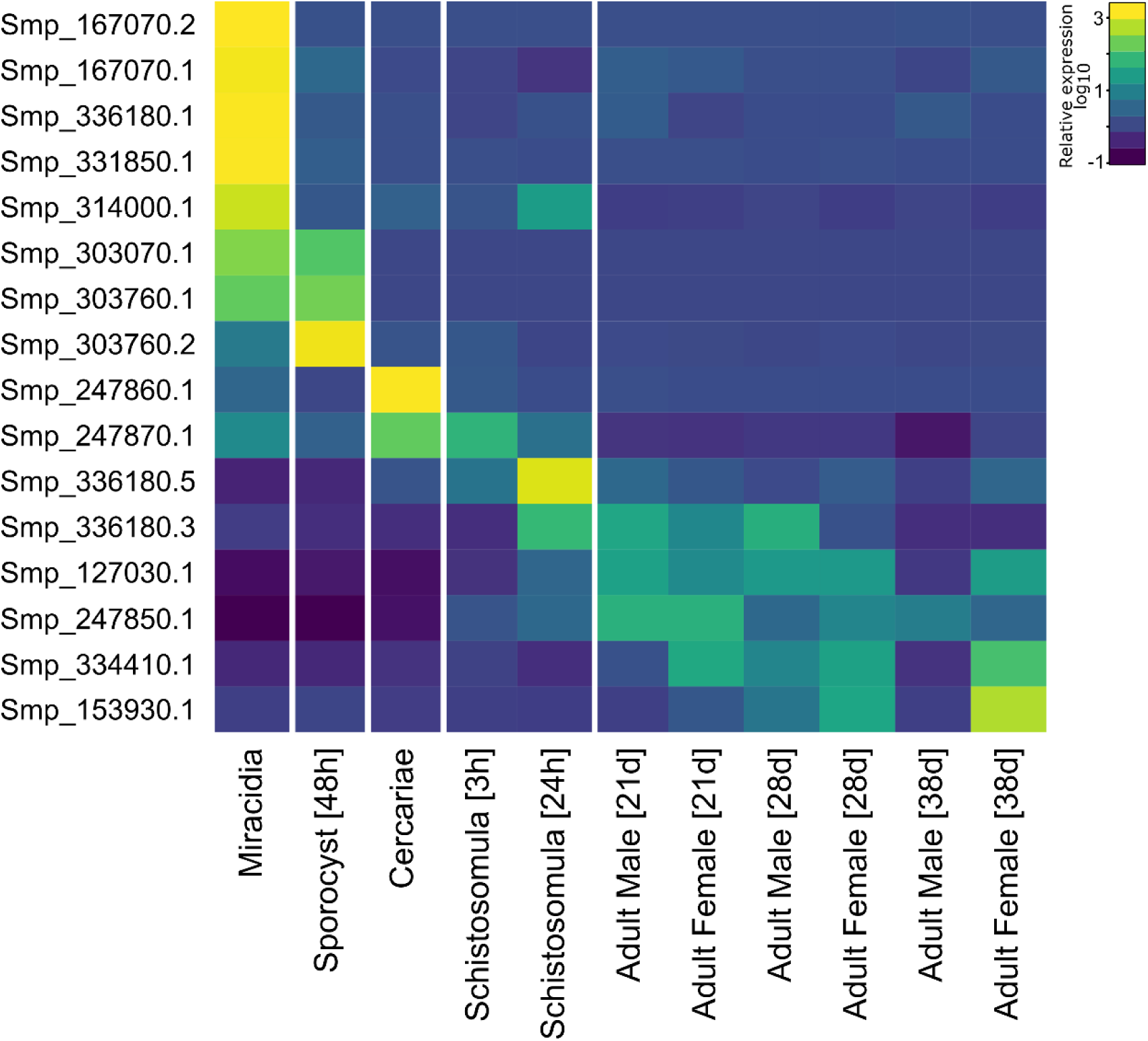
Expression of invadolysin paralogs in *S. mansoni*. Relative expression values for each transcript are shown at major life stages. Different invadolysin paralogs appear to exhibit a continuum of stage specific expression. Invadolysin (Smp_127030) is primarily expressed in *S. mansoni* adult worms. “Smp” numbers refer to the accessions for the *Schistosoma mansoni* homologs.

